# Mobile genetic elements accompany the emergence of clonal crop lineages from weed-associated populations of *Xanthomonas campestris*

**DOI:** 10.64898/2026.10.29.762866

**Authors:** Cole Sawyer, Zoë E. Dubrow, Sara C.D. Carpenter, Miles Buddy, Paul C. Owen-Smith, Adam J. Bogdanove, Christine D. Smart, Michael R. Fulcher, Matthew A. Tancos

## Abstract

*Xanthomonas campestris* is a destructive pathogen of cruciferous vegetables and ornamentals and is frequently isolated from related weed species growing near agricultural land. However, the evolutionary relationships, genomic features, and biosecurity significance of weed-associated strains remain poorly understood. Here, we integrate phylogenomic and pangenomic analyses of *X. campestris*, including nine newly sequenced strains isolated from invasive garlic mustard (*Alliaria petiolata*), to compare strains from cultivated and non-cultivated hosts across multiple pathovars. Weed-isolated strains predominantly clustered within basal, genetically diverse lineages associated with broad host ranges, whereas crop-isolated strains were concentrated in host-associated clonal lineages. The emergence of specialized crop-associated lineages coincided with multiple inferred acquisitions of large mobile elements, including integrative conjugative and mobilizable elements and plasmids carrying genes previously associated with pathogenicity and host range. Although these elements were enriched in clonal crop-associated lineages, they also occurred in diverse weed-associated strains, indicating that genetic material associated with crop infection is distributed among divergent lineages and persists in wild and weedy systems. These findings identify invasive weeds as ancillary hosts that may harbor pathogen diversity and provide opportunities for genetic exchange, underscoring the importance of surveillance beyond agricultural fields to monitor pathogen diversification and emerging threats. Our pangenomic analysis also identified crop-associated pathogen factors and putative host resistance targets for future investigation, providing a foundation for developing critically needed black rot-resistant cultivars.

**SIGNIFICANCE:** Weedy plants are often overlooked in agricultural disease surveillance, yet they can harbor pathogen diversity relevant to crop health. We analyzed 576 genomes of *Xanthomonas campestris*, including nine sequenced strains from invasive garlic mustard. Weed-isolated strains predominantly belonged to genetically diverse, broad-host-range lineages, whereas crop-isolated strains were often specialized and clonal. Crop-associated lineages showed repeated acquisition of large mobile genetic elements carrying genes associated with pathogenicity and host range. These elements also occurred in diverse weed-isolated strains, suggesting that weeds may harbor pathogen diversity and provide opportunities for exchange of traits relevant to crop infection. Our findings indicate that surveillance beyond agricultural fields can reveal pathogen diversification with biosecurity implications and inform the development of disease-resistant crops.

## INTRODUCTION

Brassicaceous crops are central to global agriculture, providing vegetables, oilseeds, forage, animal fodder, cover crops, biofumigants, and fermented and condiment products. In the United States, broccoli, cabbage, cauliflower, kale, canola, mustard, and rapeseed occupied more than 2.6 million harvested acres in 2025 and generated an estimated production value of US$2.9 billion (U.S. Department of Agriculture 2026). The four listed *Brassica oleracea* vegetables averaged approximately US$10,000 per acre, placing them among some of the most valuable US crops on a per-acre basis (U.S. Department of Agriculture 2026). Protecting these crops is critical for agricultural productivity, food systems, and economic security.

The diversity of cultivated brassicaceous crops reflects a long history of domestication, trade, and adaptation. Human selection in *B. oleracea* and *B. rapa* produced genetically related, yet morphologically distinct crops, such as kale (*B. oleracea* var. *acephala*), cauliflower (*B. oleracea* var. *botrytis*), bok choy (*B. rapa* subsp. c*hinensis*) and turnip (*B. rapa* subsp. *rapa*) (Cai et al., 2022; Cheng et al., 2016; Ma et al., 2026; Mabry et al., 2021; McAlvay et al., 2021; Qi et al., 2017). Despite their phenotypic and agronomic diversity, brassicaceous crops and related wild and weedy relatives remain susceptible to similar plant pathogens. Concerning bacterial pathogens, *Xanthomonas campestris* is one of the most damaging agents of brassicaceous crops worldwide, affecting food crops, ornamentals, and wild/weedy species (Dubrow et al. 2022; Vicente and Holub 2013).

Historically, *X. campestris* strains associated with Brassicaceae have been categorized based on host range, resulting in the creation of distinct pathovars, namely *campestris, raphani, incanae, barbareae, armoraciae,* and *abberans* (Vauterin et al., 1995). However, recent genomic and pathogenicity studies support a revised classification system comprised of five groups: *X. campestris* pv. *campestris*, pv. *incanae*, pv. *raphani*, pv. *barbareae*, and a ‘nonpathogenic’ group (Dubrow et al., 2022; Fargier et al., 2011). *Xanthomonas campestris* pv. *campestris* (*xcc*) is the most economically damaging pathovar, causing the vascular disease black rot of high-value cruciferous vegetables. Pathovar *incanae* (*xci*) causes bacterial blight, a vascular disease primarily associated with ornamentals (e.g., *Matthiola* and *Erysimum* spp) but recently documented in brassicaceous weeds as well (Dubrow et al., 2022; Tancos et al., 2022; Tancos & Frederick, 2020). Pathovar *barbareae* (*xcb*) appears relatively host-specific causing non-vascular leaf spots on the weed *Barbarea vulgaris* (yellow rocket), but recently, it has been isolated from the nonnative invasive brassicaceous weed garlic mustard (*Alliaria petiolata*) (Dubrow et al., 2022; Tancos et al., 2024). In contrast to the other pathovars, *X. campestris* pv. *raphani* (*xcr*) has a broad host range, capable of causing non-vascular leaf spots on brassicaceous and solanaceous hosts, and appearing ancestral to the other *X. campestris* pathovars based on recent comparative genomic studies (Paauw et al., 2025). The remaining group, designated ‘nonpathogenic’ (*xcnp*), lack the canonical type III secretion systems (T3SS) and frequently originate from brassicaceous weeds like *Arabidopsis* species, yet clearly remain within *X. campestris* based on high whole-genome average nucleotide identify (ANI) scores (Dubrow, 2021).

Recent field collections have revealed previously unrecognized *X. campestris* diversity among wild, feral, or weedy brassicaceous hosts. Numerous weed-associated strains exhibit broader host ranges than strains isolated from crops, suggesting that non-cultivated hosts may contribute to the maintenance and diversification of pathogen populations (Dubrow et al. 2022; Tancos et al. 2024). However, the evolutionary and epidemiological relationships between weed- and crop-associated strains remain unresolved. Earlier studies have identified diverse *X. campestris* genotypes in agricultural-associated brassicaceous weeds, but limited sampling and genomic data precluded robust inference of whether weed-associated strains contribute to black rot outbreaks or whether strains move between weeds and crops (Ignatov et al., 2007; Lange et al., 2022; Schaad, 1981). The capacity of non-cultivated brassicaceous hosts to harbor, select, or introduce novel genetic variants remains poorly understood, yet important to US agricultural biosecurity.

This knowledge gap is critically important as foreign brassicaceous weeds, such as hoary cress (*Lepidium draba*), perennial pepperweed (*Lepidium latifolium*), dyer’s woad (*Isatis tinctoria*), Sahara mustard (*Brassica tournefortii*), and garlic mustard, invade diverse agricultural landscapes (DiTomaso, 2000; Durka et al., 2005; Francis & Warwick, 2008; Gaskin et al., 2013, 2018; Rodriguez et al., 2024). Pathogen evolution is influenced by host susceptibility, density, and diversity, and as highly susceptible invasive weeds spread at monoculture levels near agricultural land, the risks of genetic exchange between different pathogen lineages increase (Goss et al., 2020; Linde et al., 2016; Penczykowski et al., 2026). Consequently, there are increased opportunities for pathogen evolution and emergence events to occur across diverse agricultural landscapes.

This study investigated the genomic diversity and evolutionary relationships of *X. campestris* strains isolated from cultivated and non-cultivated brassicaceous hosts worldwide. For simplicity, we refer to strains originating from cultivated plants as crop-isolated and those from wild, feral, or weedy plants as weed-isolated. Here, we combine nine newly sequenced *X. campestris* genomes representing *xcc, xci,* and *xcb* from invasive *A. petiolata* populations in the United States (Tancos et al. 2024) with a global collection, yielding a total of 567 genomes, to characterize the genomic features associated with crop or weed isolation, host range, and pathogen diversification. Phylogenomic and pangenomic analyses reveal that broad host range, weed-associated lineages retain an ancestral and comparatively undifferentiated genomic state, while clonal crop-associated lineages have acquired accessory genes through the repeated acquisition of large mobile elements. This led to the identification of a persistent mega-plasmid and a widespread avirulence-associated integrative conjugative element carrying multiple pathogenicity-related genes across divergent *X. campestris* lineages, including within weed-isolated strains. These discoveries support the model of specialized crop-infecting *xcc* and *xci* lineages emerging from basal, weed-associated lineages through the repeated acquisition of pathogenicity-associated mobile elements. Moreover, the presence of these crop-associated mobile elements in genetically diverse, weed-associated strains underscores the biosecurity significance of invasive weeds as possible reservoirs of pathogen diversity and potential sites of genetic exchange. These findings support surveillance of non-cultivated hosts as a means of detecting pathogen diversification before it threatens susceptible crops. Beyond its evolutionary and agricultural biosecurity implications, our pangenomic analysis of crop- and weed-isolated strains identified pathogenicity factors specific to crop infection along with candidate host resistance targets, providing a foundation for developing black rot-resistant cultivars and mitigating the devasting impact of this global disease.

## MATERIALS & METHODS

### Strain collections and genome sequencing

*Xanthomonas campestris* strains (n = 8) previously characterized from invasive garlic mustard infestations were targeted for whole genome sequencing (Tancos et al. 2024). An additional *X. campestris* strain, FDWSRU 22-193, was collected in 2022 from symptomatic garlic mustard growing along the C&O Canal in Frederick County (Point of Rocks), Maryland. Representative diseased leaves were collected, surface sterilized, and plated, according to established protocols (Tancos and Frederick 2020; Tancos et al. 2024). Yellow, mucoid bacterial colonies were individually collected and purified. The 16S rRNA sequence was used to confirm taxonomic identity, and the representative culture was stored in 20% glycerol at -80℃. A limited host range study was performed to assess pathogenicity using previously described protocols and brassicaceous hosts (Tancos et al. 2024).

High molecular weight genomic DNA was extracted from purified *X. campestris* strains (n = 9), as previously described (Tancos et al. 2022). DNA was sent to the Delaware Biotechnology Institute at the University of Delaware DNA Sequencing & Genotyping Center for quality assessment using FEMTO Pulse (Agilent, CA, USA), PacBio HiFi (Pacific Biosciences, CA, USA) library preparation, and PacBio Revio long read sequencing. The genome sequences were assembled using Flye long read assembler v.2.9.6 via Geneious Prime v. 2024.0.5 (Biomatters, Auckland, New Zealand).

### Phylogeny and population structure analysis

A total of 1,003 *X. campestris* genomes were downloaded from NCBI in April 2026. For initial clustering into pathovars, average nucleotide identity (ANI) was determined using sourmash v.4.9.4 by aligning to known reference strains from each pathovar: ATCC-33913 for *xcc*, CFBP-5824 for *xci*, 5057 for *xcr* and 5053 for *xcnp* (Supplementary Dataset 1). Strains were assigned to a given pathovar based on highest ANI score. Any strain below the threshold of 95%, relative to any pathovar representative, was dropped from the study. Species-level metadata and/or corresponding publications were reviewed manually to assign host classifications to each genome. The categorical variable termed “weed status” was used to indicate host of origin. Genomes were given a weed status of “yes” whenever a host was determined to be non-cultivated, which was defined as including all non-domesticated, feral, weedy, or otherwise wild species or individuals. Genomes were given a weed status of “no” when a host was determined to be cultivated, which was defined as including all domesticated species grown intentionally as crop or ornamental plants. If the metadata available was insufficient to make this determination, for instance a host was named only to the genus level (i.e. *Brassica* sp.), the weed status was considered “not annotated (NA)”. Pathovar *raphani* was found to only have three strains from non-cultivated hosts and was therefore dropped from the pangenome analysis, as the primary goal was to compare crop- and weed-isolated strains. Additionally, if the host metadata indicated an origin outside of Brassicaceae, the corresponding strain was dropped from the analysis. The remaining strains were further curated using Quast v.5.2.0 to prioritize genomes with 100 or fewer contigs, N50 > 50,000 bp and BUSCO v.6 requiring 95% completeness using xanthomonas_odb12 (Gurevich et al., 2013; Manni et al., 2021). The geographical source of each strain included in the pangenome analysis was identified from metadata and binned into regions: Africa, Europe, East Asia, Asia, Oceania and the Americas. The *X. campestris* population was segmented into clusters via Bayesian Analysis of Population Structure (BAPS) using fastBAPS v.1.0.8 (Tonkin-Hill et al., 2019). A set of core genes was selected by using as input the selected *X. campestris* strains and the outgroup species *X. fragariae*. Core, single-copy genes were then aligned with MAFFT v.7.505 using default parameters and trimmed with trimal v.1.5 using the -automated1 setting (Capella-Gutiérrez et al., 2009; Katoh & Standley, 2013). The resulting alignments were then used as input for IQ-TREE v.2.4.0 with 1000 ultrafast bootstraps and model selection performed with modelfinder (Minh et al., 2020). Finally, fastBAPs was performed using the core gene alignment and generated the output of IQ-TREE.

### Pangenome construction and annotation

All strains were annotated using bakta v.6.0.0 prior to pangenome construction using ppanggolin v.2.2.1 with default parameters (Gautreau et al., 2020; Schwengers et al., 2021). Estimates of pangenome openness were calculated using the ppanggolin rarefaction module with default parameters. Gene families, denoted as either persistent (>95%), shell (<95% and >5%) and cloud (<5%) were then used to calculate summary statistics for each pathovar in the R statistical environment v.4.5.1. Comparisons of gene counts within each bin (persistent, shell and cloud) were made between individual BAPS clusters and between pathovars. The Shapiro-Wilkes test of normality was performed on the distribution of gene counts for each pathovar or BAPS cluster. As all datasets were found to be non-normal, the Kruskall-Wallis test, followed by Dunn’s post hoc test were performed to compare to assess differences at the pathovar and BAPS clustering levels (Dunn, 1964; Kruskal & Wallis, 1952; Shapiro & Wilk, 1965). BAPS clusters with fewer than n = 10 strains were dropped from statistical testing.

### Multiple correspondence analysis and hierarchical clustering of principal components

The gene presence-absence output of ppanggolin was analyzed using R (R Core Team, 2025). First, strains which originated from *xcnp* were dropped from the analysis owing to a large number of unique genetic variants. Second, the ordination was limited only to the shell and cloud genome by filtering out the persistent genes. Both steps were performed to lower the number of dimensions in the data set, focusing specifically on those genes which contribute most to a BAPS while maintaining rare, but potentially important cloud variants. A presence-absence matrix was then used as input for Multiple Correspondence Analysis (MCA) from the factomineR package using default parameters v.2.16 (Lê et al., 2008). The dimension contribution values were corrected using Benzecri’s method (Benzecri, 1979). The final MCA was then used as input for Hierarchical Clustering of Principal Components (HCPC) from the factomineR package using default parameters. Extraction of the vtest metric, the measure of association between a gene and fava given cluster, was then used for plotting using gggenomes v.1.1.3 (Hackl et al., 2024). To highlight syntenic blocks, minimap2 v.2.24-r1122 was used to align genomes, and alignments were visualized with gggenomes (Li, 2018).

### Clustering of mobile elements

ICEFinder v.2 and geNomad v.1.11.1 were used to locate any putative integrative and conjugative/mobilizable elements (ICEs/IMEs), bacteriophages (phages) and plasmids in the *X. campestris* genomes (Camargo et al., 2024; Wang et al., 2024). We only collected mobilome data from high-quality genomes which were (i) listed as complete in associated NCBI metadata or (ii) contained four or fewer contigs. All elements were collected and compared using sourmash, performing an all-against-all comparison measuring Jaccard distance (Irber et al., 2024). A dissimilarity matrix was then used as input for the hclust package of R using default parameters and plotted. One cluster with a high Jaccard similarity was chosen for further analysis. High quality genomes were used as input for annoTALE v.1.5 (Grau et al., 2016). Whole genome fasta files were used as input for the predict and analyze modules. Predicted effectors were then mapped to a given gene family produced by ppanggolin. To predict the presence of type VI secretion systems (T6SS), MacSyFinder v.2.1.5 and the repository TXSSCAN were used to scan all genomes within our dataset (Abby et al., 2024; Néron et al., 2023).

### Analysis of AvrB across the *X. campestris* pangenome

The coding sequences of the *avrB* homolog found within the large ICE of *X. campestris* species was collected from the pangenome using ppanggolin (Gautreau et al., 2020). All proteins were then filtered to keep only unique isoforms using the seqkit v.2.14.0 rmdup function and aligned with MAFFT as above (Katoh & Standley, 2013; Shen et al., 2024). Alignments and logos were then plotted using the ggmsa package v.3.23 (Zhou et al., 2022). The number of times a given isoform appeared in the set of all avrB proteins was collected using seqkit rmdup (Shen et al., 2024).

## RESULTS

### Multiple *X. campestris* pathovars identified from natural garlic mustard infestations

The sequenced *X. campestris* strains from garlic mustard consisted of complete, circularized chromosomes with an average coverage of 572x (Table 1). Of the newly sequenced strains, only strains 22-193 and 21-082 had extrachromosomal plasmids. ANI scores placed the nine strains into pathovars *xcc, xci,* and *xcb*. Strain 22-193 was the only *xcc* garlic mustard representative and displayed a similar host range to *xcc* type strain ATCC 33913 with characteristic V-shaped symptoms developing on garlic mustard, cabbage (*B. oleracea* var. *capitata*), and yellow California mustard (*Guillenia flavescens*). No symptoms were observed on common stock (*Matthiola incana*) or any negative controls. The host ranges of the other eight strains were previously described (Tancos et al. 2024).

**Table 1.** Newly sequenced genomes from invasive garlic mustard (*Alliaria petiolata*).

| Strain | Pathovar* | Contig Number | Length (BP) | Coverage |
| --- | --- | --- | --- | --- |
| 22-193 | *xcc* | 3 | 5,186,788 | 545 |
| 21-080 | *xcb* | 1 | 4,984,103 | 506 |
| 21-090 | *xcb* | 1 | 5,048,307 | 596 |
| 21-091 | *xcb* | 1 | 5,057,017 | 587 |
| 21-021 | *xc**i* | 1 | 4,977,423 | 627 |
| 21-082 | *xc**i* | 2 | 5,023,488 | 575 |
| 21-086 | *xc**i* | 1 | 5,026,602 | 486 |
| 21-089 | *xc**i* | 1 | 4,934,099 | 633 |
| 21-088 | *xc**i* | 1 | 4,979,892 | 597 |

\* - pathovar designations are as follows; *xcc* – *campestris*, *xcb* – *barbareae*, *xc**i* - *incanae*

### Phylogenetic analysis of 567-member *X. campestris* pangenome denotes multiple genetic clusters composed predominantly of weed-isolated strains

To generate a robust *X. campestris* pangenome, with an emphasis on weed- and crop-isolated strains, we retained genomes with high quality metrics from diverse strains among all pathovar groups: *xcc*, *xci*, *xcb, xcr* and *xcnp.* Any strains outside the ANI cut-off of 0.95 were excluded (Supplementary Figure 1A). Pathovar *raphani* (*xcr*) was excluded as only three genomes were available from strains isolated from a weedy host (Supplementary Figure 1B). We then incorporated the nine newly sequenced strains isolated from invasive garlic mustard populations (Table 1). The final pangenome comprised 567 *X. campestris* genomes, including 65 (11.4%) strains classified as being weed-isolated.

A phylogenetic tree was generated via a core-genome multilocus sequence typing using *X. fragariae* as an outgroup (Figure 1). The phylogenetic tree condensed the strains into three large clades, representing the three pathovars *xcc*, *xci* and *xcnp*, with pathovar *xcb* nested within *xci*. We then used fastBAPs to further refine the population into distinct sub-populations (Tonkin-Hill et al., 2019). This analysis generated 12 total BAPS clusters, eight of which are within *xcc*, three within *xci* and one representing *xcnp*. BAPS clusters *xcc*-6 and *xcc*-7 were the only clusters of *xcc* predominated by weed-isolated strains. The remaining clusters of *xcc* had few to no weed-isolated strains, with the exceptions of strains WHRI-3819A and WHRI-3974A from *xcc*-2, which were isolated from wild mustard (*Sinapis arvensis*) and strain 22-193 from garlic mustard in cluster *xcc*-4 (Supplementary dataset 1).

**Figure 1.**
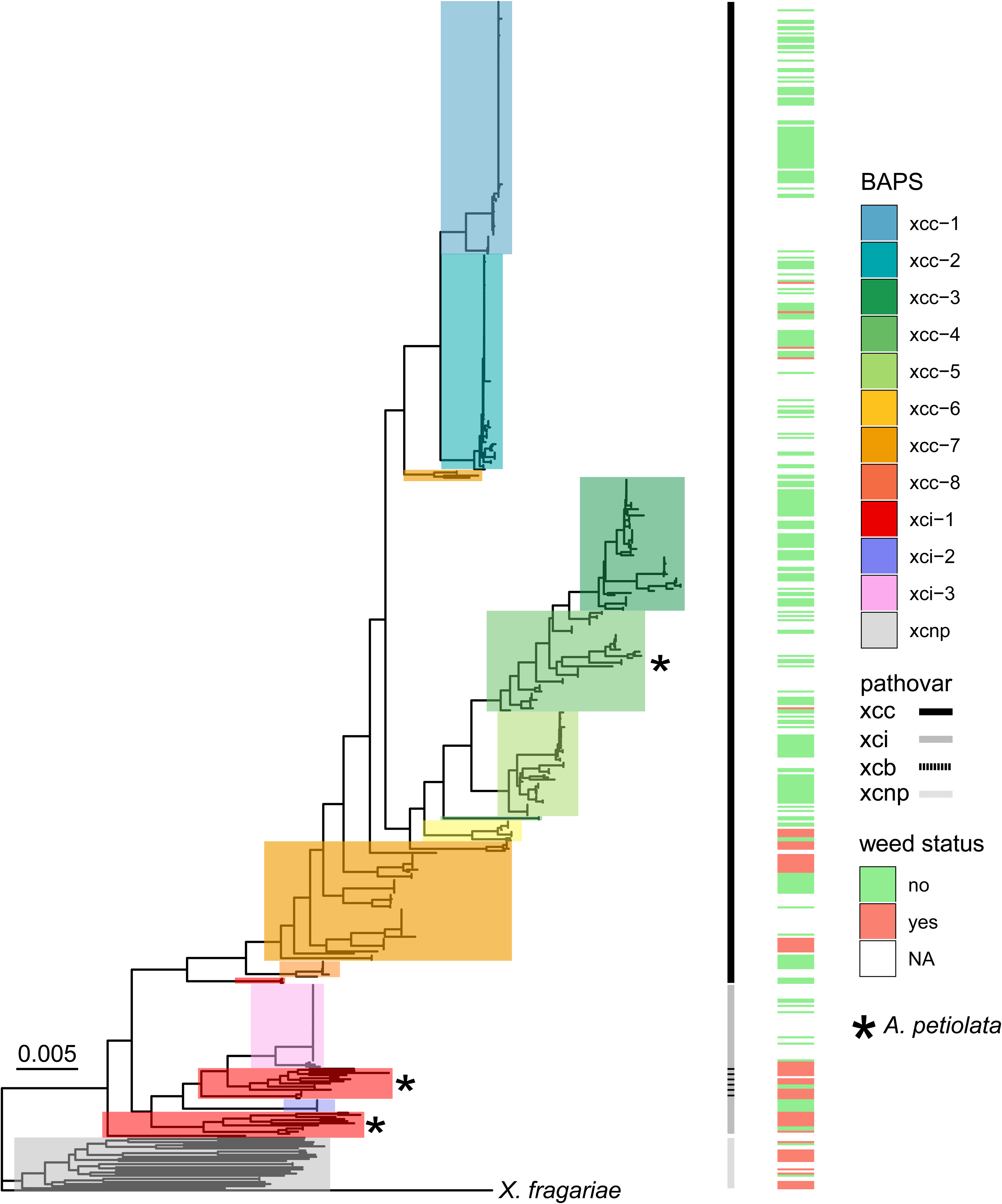
Phylogenetic analysis locates two weed-heavy genome clusters in *Xanthomonas campestris*. A) Phylogenetic tree comprising 567 genomes along with their pathovar designations (first outer ring) and weed status (second outer ring). The BAPS clustering was used to highlight the corresponding clade. BAPS clusters which contain newly sequenced strains isolated from the invasive weed *Alliaria petiolata* are denoted with a star. The branch length of outgroup *X. fragariae* was trimmed for visualization purposes.

Previous race designations of *xcc* relied primarily on host range, grouping strains by their ability to infect certain hosts (Fargier & Manceau, 2007; Singh et al., 2022; Vicente & Holub, 2013). Mapping representative strains from these races to our phylogeny suggests concordance between BAPS clusters and race designations in some circumstances (Supplementary Table 1). BAPS clusters *xcc*-1 and *xcc*-2 map to the clonal races 1 and 4 respectively, while *xcc*-6 likely represents race 6. While useful, the inclusion of race 2 within *xcc*-6, and the grouping of races 3 and 8 into a single BAPS cluster (*xcc*-3) suggests that the relationship of BAPS and races do not always correspond to each other.

BAPS cluster *xci*-1 (n = 30) and *xcnp* (n = 25) were also found to contain many weed-isolated strains, while *xci*-2 and *xci*-3 represented crop(ornamental)-isolated strains. Five of the recently sequenced garlic mustard *X. campestris* strains were located within *xci*-1 and spread between pathovar *xcb* and *xci*, demonstrating that invasive garlic mustard is a viable alternative host for *xci, xcb,* and *xcc* (Table 1). Similar to *xcc*, *xci* is expanded compared to previous phylogenetic analyses and highlights the close genetic relationship between crop- and weed-isolated strains. Specifically, a large clonal crop-infecting lineage isolated from wallflower (*Erysimum* sp.) shares recent common ancestry with garlic mustard strains and represent BAPS cluster *xci*-3. These strains largely originate from Europe, specifically the United Kingdom (Supplementary Dataset 1).

### Weed-associated genetic clusters have a larger number of isolation sources, including weed and crop species

Previous analyses of weed-isolated strains of *xcc* and *xci* noted an increased host range (Dubrow et al., 2022). Crop-isolated strains, such as reference strain ATCC-33913, could infect *B. oleracea*, wild mustard, and garlic mustard, while weed-isolated strains could additionally infect ornamentals (Dubrow et al., 2022; Tancos et al., 2024). Using metadata, we sought to delineate the relationship between genetic clustering and host (Figure 2). This analysis demonstrated that weed-associated genetic clusters have the largest number of isolation sources at the genus level, with *xcc*-6 isolated from six genera and *xci*-1 isolated from nine genera (Figure 2a). The remaining genetic clusters are far less varied in their isolation sources. Examples include genetic cluster *xcc*-1 and *xcc*-2 that are predominantly associated with *Brassica* crops.

**Figure 2.**
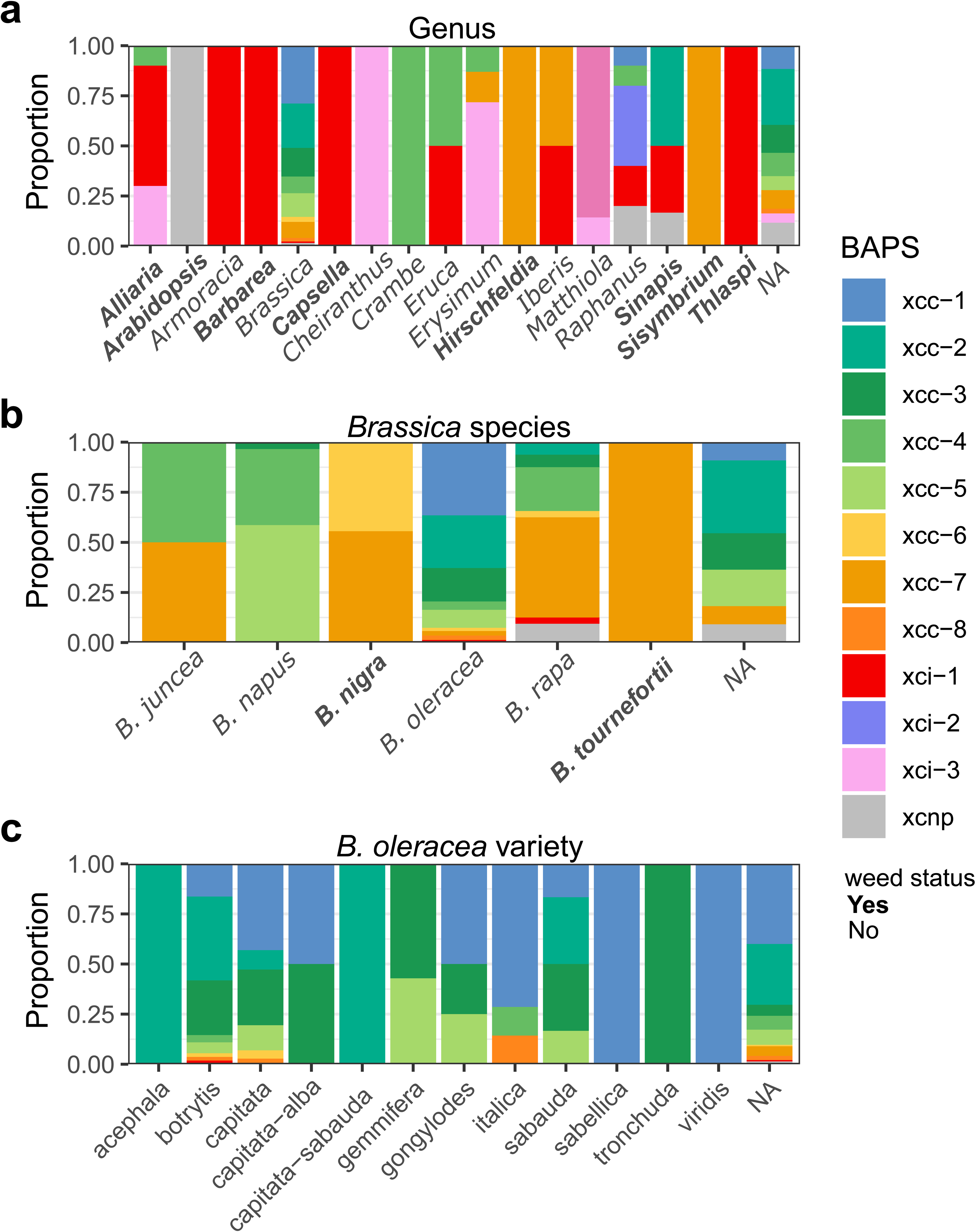
Isolation sources of BAPS clusters at varying phylogenetic levels. Each bar represents the percentage of strains that were isolated from a given genus (a) *Brassica* species (b) and cultivar of *B. oleracea* (c). Genera which contain only weed species in our analysis, as well as specific weedy species of *Brassica*, are denoted in bold.

We then sought to determine whether any associations exist between genetic clusters and the differing *Brassica* species (Figure 2b). Like the genus level, we find that *xcc*-7 has the broadest number of host species (n = 4) and is the only clade isolated from the invasive weed Sahara mustard. Cluster *xcc*-6 is the only cluster outside of *xcc*-7 which is found on the naturalized weed black mustard (*B. nigra*). The main crop species, *B. oleracea* and *B. rapa*, contain the highest diversity of genetic clusters. We note that of the two large clonal lineages *xcc*-1 and *xcc*-2, only *xcc*-2 is found on *B. rapa*. As *B. oleracea* represents the largest number of strains, we sought to determine if any relationship existed between genetic cluster and crop variety (Figure 2c). Of the varieties accurately annotated in provided metadata, cauliflower (var*. botrytis*) and cabbage (var. *capitata*) have the highest number of strains with 55 and 72, respectively. Comparing *xcc*-1 and *xcc*-2, we find that *xcc*-1 contains a higher proportion of strains found on cabbage, while the inverse is true for cauliflower.

### Analysis of the pangenome finds that weed-associated genetic cluster *xcc*-7 is associated with smaller, undifferentiated accessory genomes

The pangenome contains all genes within a representative sample of an organism, where each gene can be further classified by the percentage of strains that possess it. The persistent genome (>95% of strains) represents those genes which are core to the species and often encode critical housekeeping processes (Gautreau et al., 2020). The accessory genome is separated into two sub-categories: the shell genome (>5 to 95%) and the cloud genome (<5%) that encompass genes specific to sub-populations and genes that are rare in the pangenome, respectively. The total pangenome of Brassicaceae-isolated *X. campestris* strains is composed of 13,386 gene families, with most genes found in the cloud category (n = 7,262) and the remaining found in shell (n = 2,692) and persistent (n = 3,432) categories (Figure 3a). *Xcc* was found to have a larger shell genome on average than either *xci* or *xcnp* (Figure 3b). The cloud genome of both *xcc* and *xci* are similarly small compared to the large cloud genome of *xcnp* (Figure 3c). Separating into constituent BAPS clusters, basal lineages, such as *xcc*-7 and *xci*-1, had smaller shell genome sizes on average while the clonal lineages of *xcc*-1 and *xcc*-2 had the largest shell genomes (Figure 3d). The inverse was seen for the cloud genome, where the clonal lineages *xcc*-1, *xcc*-2 and *xci*-3 had few to no cloud genes (Figure 3e). After delineating the differences in size of the accessory genome, we sought to determine the differences in content.

**Figure 3.**
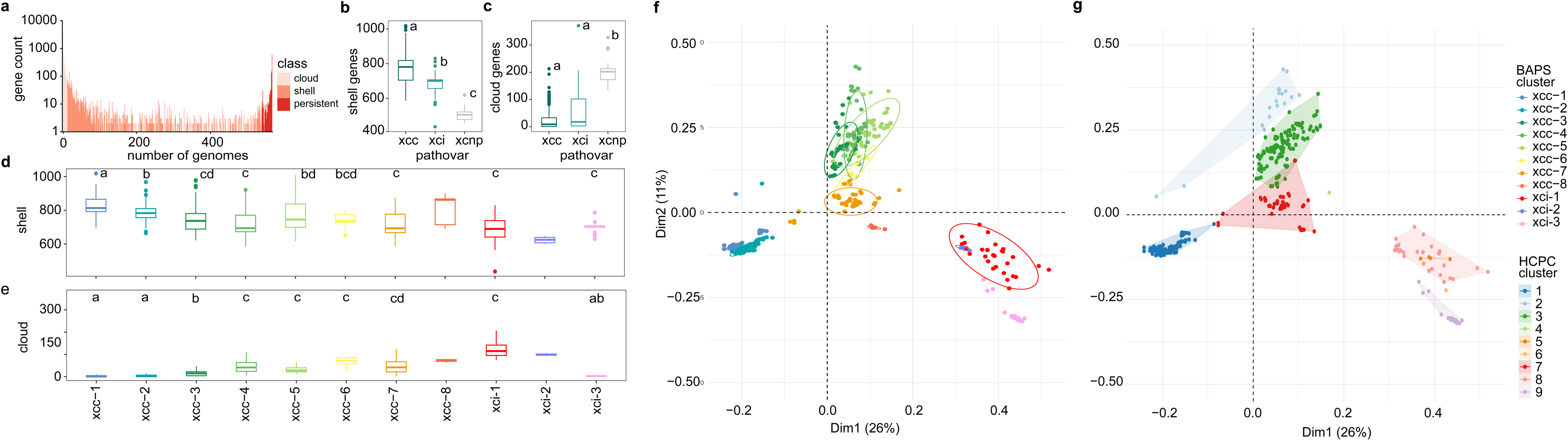
The pangenome of Brassicaceae-isolated *Xanthomonas campestris* strains. A) U-shaped plot of the pangenome, where each bar represents the number of gene counts in the number of genomes. Bars are colored by their class distinction (cloud, shell or persistent). B-C) Shell and cloud genome counts for each of the three main pathovars, *xcc* (dark blue), *xci* (blue-green) and *xcnp* (grey). D-E) Shell and cloud genome counts for individual BAPS clusters of *xcc* and *xci*. For all count comparisons, the Shaprio-Wilkes test was used to determine normality. As all datasets were denoted as non-normal, the Kruskall-Wallis test, followed by Dunn’s test, were performed to find statistically significant (P < 0.05) differences in gene count. F) MCA ordination using the accessory genome. G) Hierarchical clustering of principal components was then performed to generate clusters.

Ordination of the pangenome with MCA largely corroborated the BAPS clusters (Figure 3f) with some exceptions and outliers. The weed-associated cluster *xcc*-7 falls near the origin, highlighting a lack of strongly correlating features, except for four unique strains (MBG-39.2, WHRI-8497, WHRI-8500, CN03) which are found in between *xcc*-7 and *xcc*-2. The crop-associated clusters *xcc*-3, *xcc*-4 and *xcc*-5 are dispersed primarily along dimension two. The clonal *B. oleracea*-associated clusters *xcc*-1 and *xcc*-2 form a grouping away from the other *xcc* strains along dimension one. *Xcc*-8 is the closest BAPS cluster to *xci* while remaining next to pathovar *xcc* on the ordination, representing a basal *xcc* lineage. Additionally, multiple outliers were noted for all major clusters of *xcc*, with many extending into the positive range of dimension two. All strains of pathovar *xci*, along with three *xcc* strains (bgIFP-6745, bgIFP-6746, GBBC-3139), group away from *xcc*. While BAPS cluster *xci*-2 is closely tied to *xci*-1 in ordination space, the clonal crop(ornamental)-infecting lineage *xci*-3 has separated along dimension two, suggesting a differentiation in accessory genome content.

HCPC was then performed to determine whether relationships in accessory genome content exist when phylogenetic relationships are disregarded (Figure 3g). This analysis delineated nine clusters, five for *xcc* and four for *xci*, labeled as clusters HCPC-1 through HCPC-9. Strains from *xcc*-7 and *xcc*-6 which contain most weed strains, as well as basal lineage *xcc*-8, form HCPC-7. Strains from the crop-heavy BAPS of *xcc*-1 and *xcc*-2 form HCPC-1. The clustering failed to group the four intermediate strains between HCPC-1 and HCPC-7, splitting phylogenetically related strains between the two clusters. The crop-associated, non-clonal strains of *xcc*-3, *xcc*-4 and *xcc*-5, form HCPC-3, suggesting similar genomic content, while many of the aforementioned *xcc* outliers form their own cluster, HCPC-2. A similar trend was observed in *xci*, where most weed-isolated strains form HCPC-8, while the clonal crop-infecting lineage, and three weedy garlic mustard strains (21-088, 21-082, 21-021) form cluster HCPC-9. Additionally, multiple strains fell into clusters containing very few strains, including NCPPB-347 (HCPC-4), 11-19 (HCPC-6) and strains XUMN, 1441, I442 and I443 (HCPC-5).

### Clusters of orthologous gene (COG) family enrichment and genomic density analyses reveal association between mobile elements and differentiation of HCPC clusters

COG families are grouped by their functionality and allow for insights into the utility of a given gene set. We performed COG enrichment analysis for all HCPC clusters to determine which features are driving the differentiation between HCPC clusters. The small clusters of HCPC-4 through HCPC-6 were removed due to the low number of strains (n < 5) (Supplementary Figure 2). In all HCPC clusters, COG families representing the mobilome and extracellular structures were enriched (fold enrichment > 1). Other COG families which are enriched in multiple HCPC clusters include defense mechanisms (n = 5), intracellular trafficking (n = 4) and chromosome structure. Of the HCPC clusters, HCPC-2 (outlier *xcc* strains) had the highest fold enrichment values, with multiple COG families (cell motility, extracellular structures, chromosome structure and dynamics) above a fold enrichment of 2.

The positively associated genes for each gene cluster were then mapped onto representative strains to visualize the locations and clustering patterns of genes (Supplementary Figure 3). Given the presence of COG features associated with mobile elements, we mapped the presence of ICEs/IMEs and phages onto the genomes. For each HCPC cluster studied, we found multiple genomic islands, consisting of high densities of associated genes which align with predicted ICE insertion events or plasmids. HCPC-2 was the only cluster where most genes are mapped to a large mega-plasmid (140 – 200 kb) as seen in *xcc* garlic mustard strain 22-193 (Supplementary Figure 3b). We also note a highly associated plasmid within *xci* strain CFBP-1606 of HCPC-9 (Supplementary Figure 3h). Of the three clusters with few strains, HCPC-5 (XUMN) and HCPC-6 (11-19) both had complete genomes and were used for visualization (Supplementary Figure 3d-e). This analysis also found the presence of multiple large genomic islands in both strains that were mapped to predicted mobile elements.

### Evidence of mobile element acquisition in multiple weed-isolated strains

Given the relationship between mobile elements and the accessory genome, we sought to more fully characterize large genetic islands, focusing primarily on ICEs and plasmids because of their propensity to move large numbers of virulence-associated genes (Liu et al., 2022). To prevent misleading annotations from the use of fragmented genomes, this analysis included only complete genomes consisting of four or less contigs (n = 159 *Xc* strains). Predicted DNA sequences were then clustered following comparisons with sourmash and binned into classes, resulting in 23 classes of plasmids and 144 classes of ICE/IMEs (Supplementary Figures 4-5) (Irber et al., 2024). Given the ability of larger elements to generate high levels of variation, as seen with the case of HCPC-2, we chose to highlight the presence and sequence variation of one large plasmid and one large ICE element, beginning with the HCPC-2 mega-plasmid.

Extracting the high-quality plasmids from the dataset, we found that the mega-plasmid separated into three sub-groups, one located primarily in the United States and the others located in Canada or Taiwan (Figure 4a). All strains which contain the mega-plasmid and have metadata, apart from *xcc* garlic mustard strain 22-193, were isolated from *Brassica* crops. Further analysis of mega-plasmid annotations located multiple virulence factors including a transcription activator like-effector (TALE) shared across all variants, *xopE* in all but the Taiwanese variant, as well as a *xopJ,* and pectate lyase homolog (Figure 4b).

**Figure 4.**
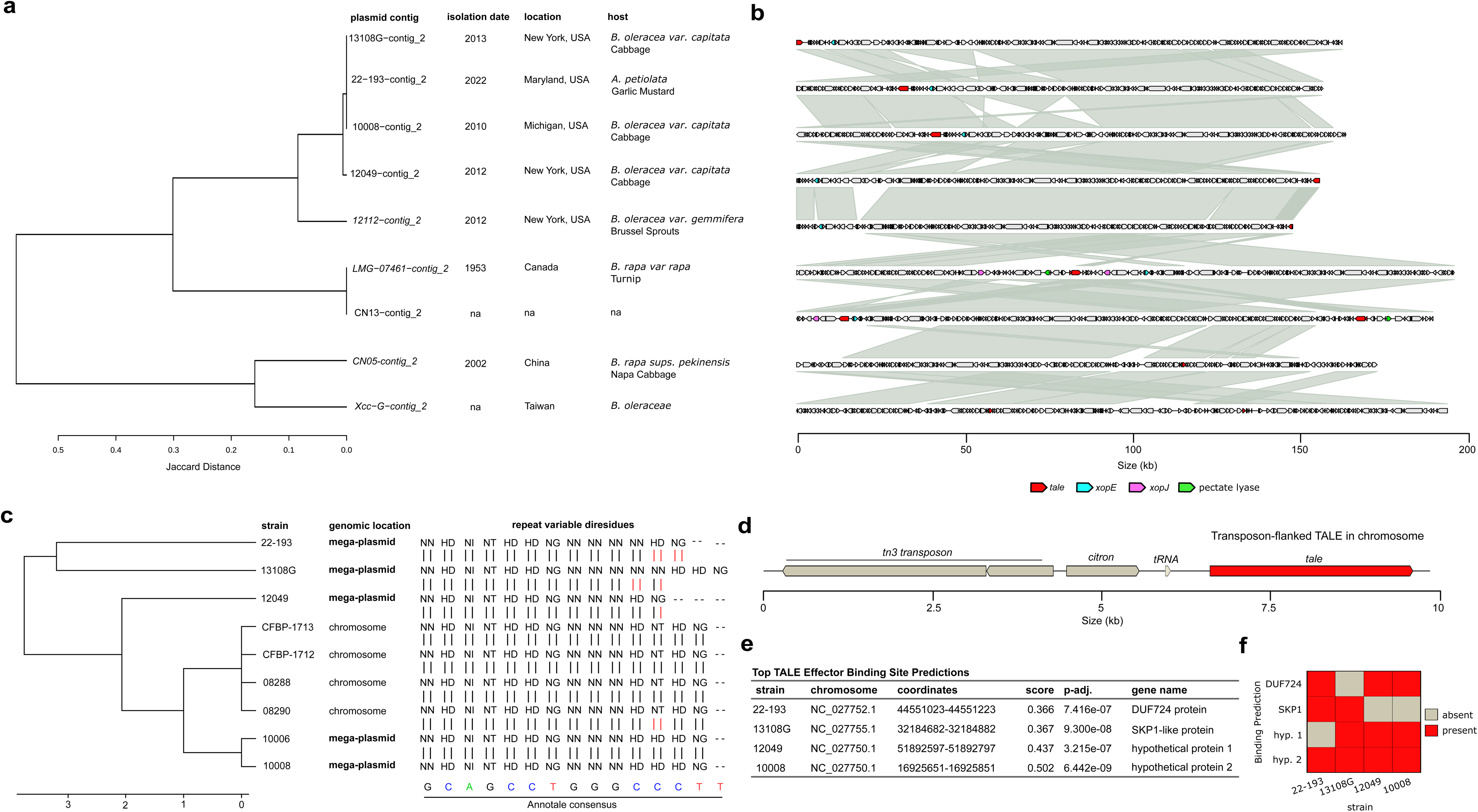
A large mega-plasmid found in a weed-isolated strain carries a novel TALE effector. A) Extraction of the hierarchical cluster from Supplementary Figure 4 containing plasmids from high-quality genomes corresponding to the mega-plasmid of HCPC-2. Isolation date, location and host (scientific and common name), if available, are supplied. B) Syntenic map of the mega-plasmid with predicted virulence associated proteins highlighted. C) Clustering analysis output by annoTALE software for the mega-plasmid TALE and related TALEs from the chromosomes of other strains. D) Genetic map of strain 08288 TALE and a nearby Tn3 transposon, as well as a linked citron protein and tRNA. E) Top binding site for each unique RVD sequence listed in sub-panel C using annotations from B. oleracea TO1000. F) Presence-absence matrix for predicted binding, where present means that binding was predicted for a given RVD and absent means that binding was not predicted for a given RVD.

The TALE was then extracted for further analysis (Figure 4c-f). All TALE proteins from the high-quality genome set were predicted and clustered to locate proteins with similar repeat variable di-residues (RVDs) using annoTALE (Grau et al., 2016). The RVDs of a given TALE determine the effector binding element (EBE) in the plant promoter which facilitates up-regulation during infection. This analysis binned the plasmid-encoded TALEs into three clusters, one set from strains isolated in the United States and the other two containing one TALE each from CN13 (Supplementary Table 2). The first cluster from the United States, including the garlic mustard isolate 22-193, was chosen for further analysis. The mega-plasmid TALE had a consensus EBE spanning 14 bp and shared a similar RVD pattern with several TALEs not found on plasmids (Figure 4c, Supplementary Table 2) (Grau et al., 2016). Several plasmids were on the edge of the contig which may affect the prediction of RVDs and was noted in the table. Comparison of this EBE with those previously described in the literature found that this class of TALEs belongs to EBE group II. The TALE proteins were re-named using standard conventions with tal13d from strain 22-193 representing a novel binding sequence.

All TALEs, both chromosomal and plasmid-borne, were found to be within proximity of a Tn3 transposon, suggesting a common route of acquisition (Figure 4d). The promoter sequences of cultivar TO1000, a reference genome of wild *B. oleracea*, was then used to search for potential binding sites of this TALE using the unique RVD sequences, represented by strains 22-193, 12108G, 12049, and 10008. This identified four proteins, a DUF724 protein, a SKP1-like protein and two hypothetical reading frames (Figure 4e). Of these, only one of the two hypothetical proteins was predicted to be bound by all four RVD sequences (Figure 4f)

Within the ICEs, we noted one large clade of highly similar elements which (i) correspond to an ICE that is strongly correlated with the clonal, crop-associated *xcc* cluster HCPC-1, (ii) contains multiple weed-isolated *xcc* strains from HCPC-7 and (iii) contains one of the ICEs in *xcb* strain 11-19, which forms HCPC-6 (Supplementary Figure 5b). To ensure collection of all ICEs in the dataset, we performed a presence-absence analysis using the gene families of the predicted ICE (Supplementary Dataset 2). This analysis also located multiple IMEs in the genomes of *xci* strains, primarily from the clonal cluster HCPC-9. From this new set of related ICEs/IMEs, a subset was collected for comparison against a strain which lacks this element, *xci* 21-089 (Figure 5). These related ICEs/IMEs were chosen for their broad phylogenetic distribution across the differing pathovars, their presence within high-quality genomes, and their putative structural variation from the presence-absence plot (Supplementary Dataset 2). This found that many of the ICEs were flanked by similar core genes, suggesting a conserved locus of integration. Comparing the gene content across these similar elements revealed the presence of one to three virulence-associated effectors, *avrB*, *avrXacE2* and *xopH*. The presence of these three effectors led us to label the element as avirulence ICE (ICE-avr). Effector *avrB* is ubiquitous throughout BAPS clusters *xcc*-1 through *xcc*-5 and is absent from all weed strains investigated, except for *xcc* weed strain 39-3. Further analysis of these strains found that the protein is truncated in 39-3 and likely nonfunctional (Supplementary Figure 6). Effectors *avrXacE2* and *xopH* are found throughout *xcc*-1 and *xcc*-2, as well as the weed-isolated strains with some structural variation, as can be seen with *avrXacE2* in *xcc*-2. Additionally, we noted multiple operons which appear to serve unknown/non-pathogenic functionalities, including coproporphyrinogen synthesis, an operon containing biosynthetic proteins, methyltransferases, several DNA modification modules and a large T6SS unique to the weed-isolated *xcb* strain 11-19 (Figure 5). To determine whether other *X. campestris* strains contain a T6SS, we utilized MacSyFinder to search the pangenome (Abby et al., 2024). This analysis found that many isolates from *xcnp* contain a T6SS, but *xcb* 11-19 is the only pathogenic strain with this same class of system (Supplementary Table 3). The uniqueness in size and content of the T6SS in 11-19, relative to *xcnp,* led us to search for similar ICEs in other *Xanthomonas* species. A protein blast against the nr database of NCBI located homologous ICEs in strains of the species *X. hortorum* which also contain a homologous T6SS.

**Figure 5.**
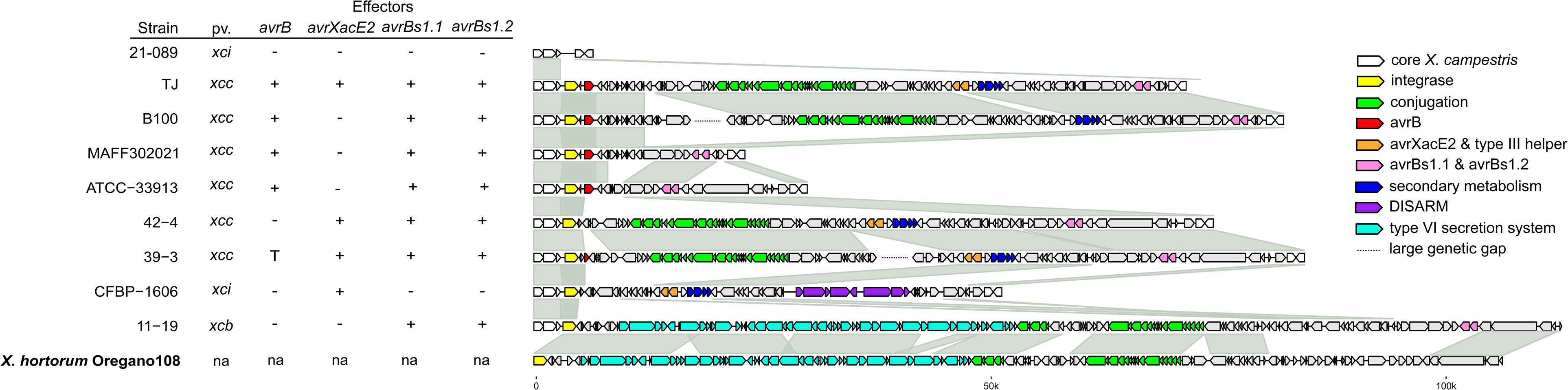
A large, effector-carrying ICE has integrated into weed-isolated *Xanthomonas campestris* genomes. Representative strains from the clustering analysis in Supplementary Figure 5 chosen to demonstrate the variety of structural variations and genetic features associated with ICE-avr. The pathovar designation as well as the presence of a given effector protein (+, present; -, absent; T, truncated, na, not available). Genetic maps of the corresponding loci for each ICE are also displayed. Large genetic gaps (> 20 kb) are highlighted to denote putative DNA inversions which have split portions of the ICE to different locations in the genome. Bold represents a strains from outside *X. campestris*, highlighting a potential horizontal gene transfer event.

## DISCUSSION

Genome sequencing of nine additional *X. campestris* strains isolated from the invasive weed garlic mustard expanded the number of well characterized weed-associated strains available for comparative genomic analyses. These data are critical for understanding the role of wild alternative hosts in bacterial evolution and the potential transfer of virulence-associated genetic elements among strains. Although several studies have shown that taxonomically related wild hosts can harbor diverse *X. campestris* populations (Dubrow et al. 2022; Ignatov et al. 2007; Krauthausen et al. 2018; Lange et al. 2022; Tancos et al. 2024), this is the first study to (i) identify the genetic features that distinguish crop- and weed-isolated *X. campestris* strains, including the acquisition and retention of pathogenicity related mobile elements that characterize crop-associated lineages, and (ii) document the presence of these large mobile elements within diverse weed-associated lineages. We hypothesize that the acquisition of mobile elements contributed to the selection and emergence of highly clonal, crop-infecting lineages in both *xcc* and *xci* that proliferated as high-yielding brassicaceous crops were cultivated, selected, and traded over human history. From a biosecurity perspective, the recurrent identification of these elements in basal lineages highlights the role of weeds as possible reservoirs and potential ancillary hosts for genetically diverse *X. campestris* populations. By harboring strains capable of carrying mobile genetic elements implicated in crop pathogenicity, weeds may provide opportunities for the emergence, persistence, and dissemination of new pathogen lineages with enhanced virulence potential or varied host ranges.

### Phylogenetic and isolation source data suggests that basal lineages of X. campestris, regardless of pathovar, have increased host ranges

Phylogenomic and BAPS cluster analysis provided new insight into the host range and evolutionary history of both *xcc* and *xci.* Historically, *xcc* has been separated between six and nine races based on host competency (Vicente & Holub, 2013), compared to the eight BAPS we defined. Interestingly, x*cc* race 6 has the largest number of viable *Brassica* host species (Vicente & Holub, 2013), and is represented in BAPs cluster *xcc-*7 with race-characterized strain WHRI-6181. Prior work from our group has demonstrated that many *xcc*-7 strains, previously labeled as *xcc*-CA (California), infected more species than their crop-isolated counterparts and comprised a distinct group found in both California and China (Dubrow et al., 2022). Our analysis of host isolation source corroborates these studies and suggests that the basal clade *xcc*-7 has a broader host range when compared to crop-associated lineages. Our study also placed multiple new strains in this clade, expanding its geographic range into Europe, primarily England and Portugal. Many of these European strains were found to infect either wallflower (*Erysimum*) or *Brassica* species. While limited publicly available metadata prevents definitive classification of these strains as either weed or crop-associated, their isolation from those genera suggests that weed-associated strains of *xcc*-7 may be able to infect important crop species and/or ancestral landraces.

The widespread global distribution of strains within the basal *xcc*-7 clade may reflect historical movement of *Brassica* germplasm. Historically, Portugal, China, and California share relevant agricultural trade and human migration links that could provide a plausible explanation for this geographic range. Being in Mediterranean Europe, Portugal contains unique, ancient *Brassica* landraces in the region where *B. oleracea* domestication is thought to have occurred, and was the likely source of germplasm introduced into China by the fifth or sixth century, contributing to the development of unique Chinese cultivars, such as Chinese kale (Cai et al., 2022; Dias, 1995; Luo et al., 2025; Mabry et al., 2021; Zhang, 2009). Later, Portugal’s extensive maritime empire with China in the sixteenth century may have further facilitated the movement of plant material between these regions (Casimiro, 2020). In the nineteenth century, mass migration from China to California may have provided a major pathway for the introduction of Asian *Brassica* species into the western United States (Galenson, 1987; Molinar, 2012; Rollins & Al-Shehbaz, 1986). Although contemporary observations and records of diversity in brassicaceous hosts and *X. campestris* populations are consistent with such a scenario, it remains speculative and could be further evidenced through additional genomic, historical, and population-level analyses.

Crop-associated strains of *xcc* can be separated into six clusters, including two clonal lineages (*xcc*-1 and *xcc*-2) associated with cabbage and cauliflower infections. *Xcc*-1 and *xcc*-2 were found to represent *xcc* races 1 and 4, the major sources of brassicaceous crop infections globally (Vicente & Holub, 2013). Analysis of host-isolation source metadata found that these clades had strict host ranges, often infecting only crop varieties of *B. oleracea*. Additionally, we noted very few instances of weed-isolated strains coming from crop-associated BAPS clusters. This finding could reflect (i) sampling biases as few studies investigate weedy species, (ii) the limited or unidirectional movement of pathogen strains between crops and weeds, or (iii) genetic factors that limit weed-associated strain competitiveness on widely cultivated crop species. Although naturalized weeds do not appear to serve as primary inoculum sources for major black rot outbreaks in New York or California (Ignatov et al., 2007; Lange et al., 2022; Schaad, 1981), the potential exchange of genetic elements between crop- and weed-isolated strains remains unclear because the previous studies used multilocus sequence analysis or fingerprinting methods that could not detect such exchanges. Moreover, the movement of strains between weeds and crops may simply be infrequent or confined to unique geographic regions that contain highly conducive host community landscapes or environmental features. Future research should focus on isolating more strains of *xcc* from weedy hosts across different landscapes and determining their relationship to proximate crops.

In contrast to *xcc* which is predominately isolated from crop hosts, *xci* is isolated from similar numbers of crop and weed host species. While the pathovar is named after its traditional host, common stock, analysis of the isolation source of these strains suggests that *xci* has the broadest host range of the Brassicaceae-isolated *X. campestris* pathovars. This comprises many weed species, including shepherd’s purse (*Capsella* sp.), pennycress (*Thlaspi arvense*) and invasive garlic mustard. BAPS clustering denoted three different genetic clusters, the polyphyletic *xci*-1, a small clonal lineage of *xci*-2 and a large clonal crop-infecting lineage of *xci*-3. *Xci*-1 is remarkable in that it contains several strains from pathovar *xcc* within its cluster.

Given that both the ANI clustering algorithm used to assign pathovar designations and the core-genome MLST assign these strains to *xcc*, the inclusion of these strains within *xci*-1 suggests that they may represent the lineage which first diverged from *xci* in host-preference toward cruciferous vegetables. Of the three strains of *xcc* identified in *xci-1*, two are from arugula (*Eruca sativa*), which is another Mediterranean native that has been consumed since ancient times (Song et al. 2025). Unlike related *Brassica* crops, arugula remains a relatively ‘wild’ cultivated Brassicaceae as artificial breeding and selection are challenging and comparative genomics do not support evidence of strong artificial selection (Song et al. 2025). Additionally, we note that the majority of *xci* strains are isolated from wallflower (*Erysimum* sp.), specifically a clonal lineage originating from Europe which forms its own BAPS cluster, *xci-*3. The existence of clonal crop-infecting lineages in both *xcc* and *xci* suggests selection for single, highly virulent, crop-specific genotypes across both pathovars. Given this, we sought to determine the genetic features which distinguish the weedy and crop-associated clonal lineages using a pangenomic approach.

### Accessory genome content of weed-isolated strains represents an undifferentiated starting point for the emergence of clonal, crop-associated lineages

In contrast to BAPS that represented core genes, MCA and HCPC were used to generate clusters via their accessory genomes, grouping strains together based on shared genetic features and then analyzed using COG enrichment and genomic location. Given the high number of cloud genes within *xcnp*, we excluded them from the analysis. Ordination analyses of accessory genomic content successfully differentiated *xci* and *xcc,* as well as their respective crop- and weed-associated lineages. The identification of distinct *xci* clusters, together with the phylogenomic data, suggests that a shift in accessory genome content accompanied the emergence of the clonal, crop(ornamental)-infecting *xci* lineage. This shift may have occurred in the most recent common ancestor shared by the crop-infecting lineage and the strains isolated from weedy garlic mustard. COG enrichment and genomic location data suggests that the introduction of multiple mobile elements, in the form of ICE/IMEs and phages are strong contributors to this differentiation. Of the functionalities enriched in *xci* clonal crop-associated (HCPC-9) versus weed-associated (HCPC-8), we find that HCPC-9 contains more genes associated with coenzyme metabolism and secondary metabolism, which could be beneficial *in planta*. As for smaller clusters, HCPC-6 and HCPC-5, which represent only one and three *xci* strains, respectively, we demonstrate that their placement is owed to the presence of distinct mobile elements, including two large genomic islands in *xcb* strain 11-19. This suggests that in the history of *xci*, mobile elements are a primary driver of differentiation in accessory genome content.

Ordination of *xcc* strains suggested that (i) the weedy strains (HCPC-7) represent an undifferentiated state with few genetic features delineating them and (ii) two different sets of genomic features exist for crop-associated *xcc* lineages (HCPC-1: clonal and HCPC-3: non-clonal). The finding that the clonal, crop-associated lineages *xcc*-1 and *xcc*-2 (HCPC-1) contain more shell genes on average than the *xcc* weed-associated lineage HCPC-7 suggests that the evolutionary history of the clonal, crop-associated lineage is predominantly one of gene gain. Furthermore, the placement of several unique *xcc* strains between the crop-associated cluster HCPC-1 and weed-associated cluster HCPC-7 in the ordination suggests a stepwise manner of gene gain over evolutionary time. Corroborating this finding, these unique strains also sit between the weed- and crop-associated strains on the phylogenetic tree. The evidence suggests multiple acquisitions of mobile genomic islands coincided with the emergence of the clonal, crop-associated strains of HCPC-1 from a weedy, basal genomic background (HCPC-7). A similar phenomenon occurred in multiple outlier strains of the ordination (*xcc* cluster HCPC-2), providing corroborating evidence that individual, large mobile elements strongly shape the accessory genome of *X. campestris*. *Xcc* cluster HCPC-2 is composed of multiple strains from varying BAPs clusters, including the garlic mustard-isolated strain 22-193. Analysis of the location of these genes identified a single, large mega-plasmid found in all strains of HCPC-2.

Much has been written on the association between mobile elements, the accessory genome and corresponding shifts in phenotype. This phenomenon is relatively common in bacterial populations and leads to many economically important phenotypes, including the emergence of antibiotic resistance and host-range shifts leading to emergent diseases (Ebmeyer et al., 2021; Maddock & Hulin, 2025; Watanabe et al., 2025). Examples of host range shifts can be seen in the plant pathogen *Pseudomonas syringae*, where the movement of large ICEs have led to the emergence of novel strains on multiple plant species including common bean, watermelon, cantaloupe and squash (Merfa et al., 2026; Pitman et al., 2005). The movement of these large mobile elements has been reported broadly across Gammaproteobacteria; however, little work has focused on the acquisition of these elements between sub-populations of the same species or the movement of pathogenicity effectors between crop- and weed-associated clades (Watanabe et al., 2025). Additionally, our analysis corroborates previous research on *xcr,* which is the only *X. campestris* pathovar that still possesses a CRISPR genome defense system (Paauw et al., 2025). Their work demonstrated that loss of the CRISPR system in *xcc* and *xci* allowed for the uptake of novel effectors via mobile elements. Our research combines and expands upon these findings, linking acquisition of mobile elements to shifts in host-range and highlighting the threats of a potential emergence event due to these movements.

### Evidence for multiple movements and introgressions of large mobile elements carrying multiple effectors into weed-isolated X. campestris strains

Our study suggests that mobile elements are critical to the differentiation of the accessory genome of *X. campestris*, the acquisition of which coincides with the emergence of clonal, crop-infecting lineages in pathovars *xcc* and *xci*. We also note in the case of HCPC-2, the presence of a large mega-plasmid within the garlic mustard *xcc* strain 22-193. The mega-plasmid was conserved across 10 *xcc* strains within the high-quality genome dataset representing strains from the US, Canada, and Taiwan. Despite being a rare, mega-plasmid, it was identified in both one of the oldest (LMG-07461; isolated in 1953) and newest (22-193; isolated in 2022) strains, highlighting its persistence in both crop- and weed-isolated *xcc* strains. When looking at the entire global *X. campestris* dataset, including fragmented genomes, evidence of the mega-plasmid remains rare and specific to *xcc* with an additional nine strains identified.

The mega-plasmid from US-isolated *xcc* strains is smaller in size and lacks several pathogenicity-associated genes, such as *xopJ* and a pectate lyase. However, all mega-plasmid variants share a TALE. Similar TALEs were identified on the chromosome of strains that did not contain the mega-plasmid, but the TALE was physically linked to a Tn3 transposon in both mega-plasmids and chromosomes. TALEs are often co-localized with mobile elements, so this finding is unsurprising, however, the ramifications of the TALEs presence is currently unknown and requires further study (Ferreira et al., 2015). Preliminary analysis using the predicted promoter sequences of *B. oleracea* TO1000 suggests multiple potential protein targets, but this again requires molecular validation. Previous research has demonstrated that the TALE may act through up-regulation of a berberine bridge enzyme in the *Brassica* genome, however, it is currently unknown whether the variation seen in the novel TALE tal13d found in weed-isolated strain 22-193 has similar effects *in planta* (Dubrow, 2021).

The large number of predicted ICEs/IMEs in the *X. campestris* pangenome led to the identification of a large mobile element (>100kb), ICE1, found in clonal, crop-associated *xcc* strains (HCPC-1), a few *xcc* weed-associated strains (HCPC-7), and the *xcb* strain 11-19. Presence-absence analysis with the genes from ICE1 located more potential mobile elements not identified by ICEfinder within the clonal, crop-associated *xci* strains (HCPC-9) (Wang et al., 2024) (Supplementary Dataset 2). Interestingly, the presence of a novel T6SS carried by an ICE1 variant in strain *xcb* strain 11-19 was unexpected as only *xcnp* appears to possess a T6SS in *X. campestris*. The *xcb* T6SS appears unique relative to the known *xcnp* variant. While not exhaustive, the lack of evidence for this T6SS in other strains of pathogenic *X. campestris* genomes, as well as a similar system in the pathogen *X. hortorum*, suggests that this element originated from outside of *X. campestris* and is the product of a horizontal gene transfer event from a non-*X. campestris* species. The T6SS of plant-associated bacteria have often been described as horizontal gene transfers and the placement of T6SS within ICEs have been reported in bacteria from the order Bacteroidales; however, to our knowledge, this is the first report of a horizontal gain of a T6SS in *Xanthomonas* (Bernal et al., 2018; Coyne et al., 2014). The origin of this element and its presence within other Xanthomonadales remains to be studied.

The location of the large mobile element, ICE1, was previously noted by He et al. who were the first to show a large genomic island conserved across multiple Chinese *xcc* strains (He et al., 2007). Our analysis builds on this finding, demonstrating that the ICE is widespread across global *xcc* populations and can carry upwards of three avirulence genes. These genes include *avrB*, *avrXacE2* and the type II helper protein MltB, and two copies of *avrbs1*. Given the large number of avirulence genes, we have opted to name the element the avirulence ICE (ICE-avr). *Xcc* strains from the clonal, crop-associated lineage (HCPC-1) often contain all three sets of avirulence genes, however, one lineage in the *xcc*-2 BAPS cluster appears to have lost *avrXacE2*. Of the effector genes, *avrB* is found almost exclusively in *B. oleracea*-isolated strains. Strains from other clusters have additionally lost the conjugation machinery, leading to the creation of IME-avr, and being incapable of movement outside the cell.

Weed-associated *xcc* strains 33-1, 42-1, 42-2 and 42-4 (HCPC-7) possess ICE-avr, but lack the *avrB* effector. In contrast, the Sahara mustard (invasive weed)-isolated strain 39-3 has ICE-avr as well, but contains a truncated version of *avrB* with an insertion element in the middle of the gene, disrupting the C-terminal region, and likely function. As *avrB* is highly conserved across *B. oleracea*-isolated strains, but is missing in all weed-isolated *xcc* strains, it suggests that *avrB* may be selected against in wild/weed-associated *xcc* populations owing to potential gene-for-gene interactions in weedy plant species which do not appear in domesticated *B. oleracea* crops. This is supported by a 2005 study that targeted eight *xcc* effectors for mutagenesis in type strain ATCC-33913, in which *avrB* (syn: *avrXccFM; avrXccC*) was the only phenotypically functional effector identified, conferring race-specific avirulence and expanding host range to include Florida mustard (*B. juncea*) when disrupted (Castañeda et al., 2005)

The strong association of *avrB* among crop-infecting *xcc* strains, and the confirmed molecular role of *avrB* in altering host range, suggests this avirulence protein confers a selective advantage to *B. oleracea*-infecting *xcc* strains but not weed-infecting strains, and potentially serves as an ideal black rot resistance breeding target. Based on the gene-for-gene theory, Florida mustard should carry resistance genes to *avrB* and would be a primary target for *xcc* resistance breeding (Castañeda et al., 2005; Shaw et al., 2021). AvrB was one of the first T3SS effectors investigated in *P. syringae* and is recognized by the disease resistance gene, *RPM1*, via the rhamnosylation of decoy RPM1-interacting protein 4 (RIN4), triggering the hypersensitive response, which is a programmed cell death response to prevent systemic infection (Eitas et al. 2008; Gopalan et al. 1996; Peng et al. 2024). While domesticated *B. oleracea* genomes contain annotated *RPM1*-like nucleotide-binding leucine-rich (NLR) resistance genes; to our knowledge, no published studies have functionally validated a *B. oleracea RPM1* resistance gene in domesticated varieties. This is partly because *Brassica* genomes are mesopolyploid, structurally variable, and rich in rapidly evolving NLR clusters, and annotation alone is insufficient to establish a true functional *RPM1* ortholog (Eitas et al., 2008; Gopalan et al., 1996; Peng et al., 2024). The functional validation and introgression (or transgenic transfer) of the Florida mustard *RPM1* locus should be prioritized as a strategy for breeding black rot-resistant *B. oleracea* cultivars.

Our pangenomic analysis demonstrates that early-branching *X. campestris* strains, which represent the undifferentiated starting point for *xcc* and *xci*, are primarily responsible for the infection of weedy species. These strains are globally distributed and can infect multiple weed and crop species, including the invasive and widespread weed, garlic mustard, which is the only documented host to naturally support *X. campestris* strains from three different pathovars. This highlights a critical need to monitor weedy or feral species related to high value crops because they harbor genetically diverse, broad-host range strains with the potential to initiate disease emergence events. Additionally, we demonstrate that genetic differentiation of *X. campestris* is largely driven by the movement and structural variation of large mobile elements. We highlighted two large elements, a mega-plasmid and ICE-avr, which contain virulence genes that may contribute to pathogenicity in diverse *X. campestris* populations. The successful integration of the mega-plasmid and ICE-avr element into divergent weed-associated strains further highlight the biosecurity risks of weedy pathogen reservoirs and their potential role in facilitating the global exchange of genetic material across *X. campestris* strains, pathovars, and related species. While our study focuses on *X. campestris,* it demonstrates how surveillance of weed-associated pathogen diversity, combined with pangenomic analysis, can reveal both biosecurity risks and promising resistance gene targets, informing urgently needed strategies for mitigating and managing devastating plant diseases.

## DATA AVAILABILITY

The newly sequenced garlic mustard-isolated *X. campestris* genome assemblies and annotations are made available through the USDA Ag Data Commons repository. These data are currently accessible a with temporary, private link and can be retrieved through the Figshare platform (https://figshare.com/s/18f135aa122d8bcab958).

## CODE AVAILABILITY

No custom scripts were employed in this study. All analyses were conducted in accordance with established protocols and strictly followed the standard command-line instructions provided in the official documentation of each software package.

## ACKNOWLEDGMENTS

We thank Jami L. Thomas for performing HMW *X. campestris* gDNA extractions and Brewster F. Kingham and Olga Shevchenko at the University of Delaware DNA Sequencing & Genotyping Center for their valued assistance. Funding for this research was supported by the United States Department of Agriculture (USDA), Agricultural Research Service (ARS) through project 8044-30400-001-000-D, Animal and Plant Health Inspection Service through project 8044-22000-047-022-I, and in part by an appointment of C. Sawyer to the ARS Research Participation Program administered by the Oak Ridge Institute for Science and Education through the interagency agreement between the U.S. Department of Energy and the USDA. ORISE is managed by ORAU under DOE contract number DE-SC0014664. Portions of this research were conducted under National Park Service Research Permit Number MONO-2021-SCI-0008. Portions of this manuscript text were edited with the assistance of large-language models implemented though USAi (https://chat.usda.usai.gov). The findings and conclusions in this publication are those of the authors and should not be construed to represent any official USDA, DOE, or U.S. Government determination or policy. The mention of trade names or commercial products in this publication is solely for the purpose of providing specific information and does not imply recommendation or endorsement by the U.S. Department of Agriculture. USDA is an equal opportunity provider and employer.

## SUPPLEMENTARY TABLE LEGENDS

**Supplementary Table 1.** BAPS cluster association with historical race designations.

| **Representative** | **Race** | **BAPS** |
| --- | --- | --- |
| WHRI-3811<sup>*</sup> | 1 | *xcc-2* |
| WHRI-3849a<sup>*</sup> | 2 | *xcc-6* |
| WHRI-5212<sup>*</sup> | 3 | *xcc-3* |
| *xcc-c7*<sup>†</sup> | 4 | *xcc-1* |
| CFBP-6865<sup>*</sup> | 5 | *xcc-5* |
| WHRI-6181<sup>*</sup> | 6 | *xcc-6* |
| CFBP-1124<sup>‡</sup> | 8 | *xcc-3* |

<sup>\*</sup> - Vicente et al. (2012)
<sup>†</sup> - Singh et al. (2022)
<sup>‡</sup> - Fargier et al. (2007)

**Supplementary Table 2.** Predicted unique TALEs of EBE group 2 from *Xanthomonas campestris* mega-plasmid and related chromosomal proteins.

| Strain | TALE name<sup>†</sup> | Edge of contig? | Has plasmid? | Repeat-variable diresidue | Distance to Tn3 Transposon (kb) |
| --- | --- | --- | --- | --- | --- |
| 22-193 | tal13d<sup>‡</sup> | No | Yes | NN HD NI NT HD HD NG NN NN NN NN HD NG | 16 |
| 13108G | tal15l | Yes | Yes | NN HD NI NT HD HD NG NN NN NN NN NN HD HD NG | 16 |
| 10008,10006 | tal14a | No | Yes | NN HD NI NT HD HD NG NN NN NN HD HD HD NG | 14 |
| 12049 | tal11e | Yes | Yes | NN HD NI NT HD HD NG NN NN NN HD NG | 16 |
| 08288,08290,CFBP-1712,CFBP-1713 | tal14b | No | No | NN HD NI NT HD HD NG NN NN NN HD NT HD NG | 2 |
| CN16* | tal14c | NA | No | NN HD NI NT HD HD NG NN HD NN NN HD HD NG | NA |

\* - Denotes isolates which were not in the high-quality genome set but had previously described TALEs from Denance et. al 2018
† - TALE names continued from Denance et. al 2018 following standard conventions
‡ - First description of TALE
NA – Analysis not performed

**Supplementary Table 3.**
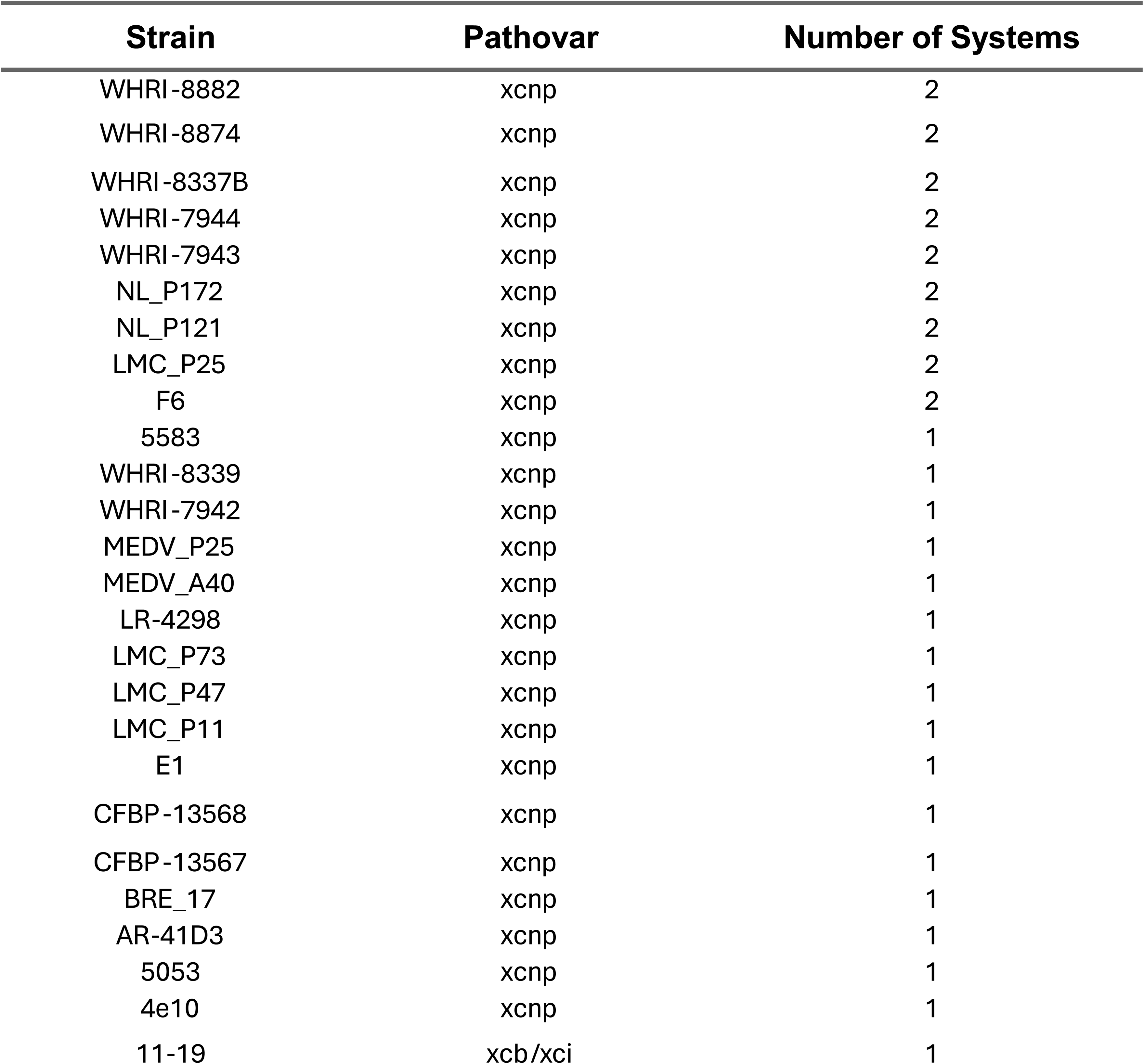
Predicted T6SS from the *Xanthomonas campestris* pangenome.

| Strain | Pathovar | Number of Systems |
| --- | --- | --- |
| WHRI-8882 | xcnp | 2 |
| WHRI-8874 | xcnp | 2 |
| WHRI-8337B | xcnp | 2 |
| WHRI-7944 | xcnp | 2 |
| WHRI-7943 | xcnp | 2 |
| NL_P172 | xcnp | 2 |
| NL_P121 | xcnp | 2 |
| LMC_P25 | xcnp | 2 |
| F6 | xcnp | 2 |
| 5583 | xcnp | 1 |
| WHRI-8339 | xcnp | 1 |
| WHRI-7942 | xcnp | 1 |
| MEDV_P25 | xcnp | 1 |
| MEDV_A40 | xcnp | 1 |
| LR-4298 | xcnp | 1 |
| LMC_P73 | xcnp | 1 |
| LMC_P47 | xcnp | 1 |
| LMC_P11 | xcnp | 1 |
| E1 | xcnp | 1 |
| CFBP-13568 | xcnp | 1 |
| CFBP-13567 | xcnp | 1 |
| BRE_17 | xcnp | 1 |
| AR-41D3 | xcnp | 1 |
| 5053 | xcnp | 1 |
| 4e10 | xcnp | 1 |
| 11-19 | xcb/xci | 1 |

## SUPPLEMENTARY FIGURE LEGENDS

**Supplementary Figure 1.**
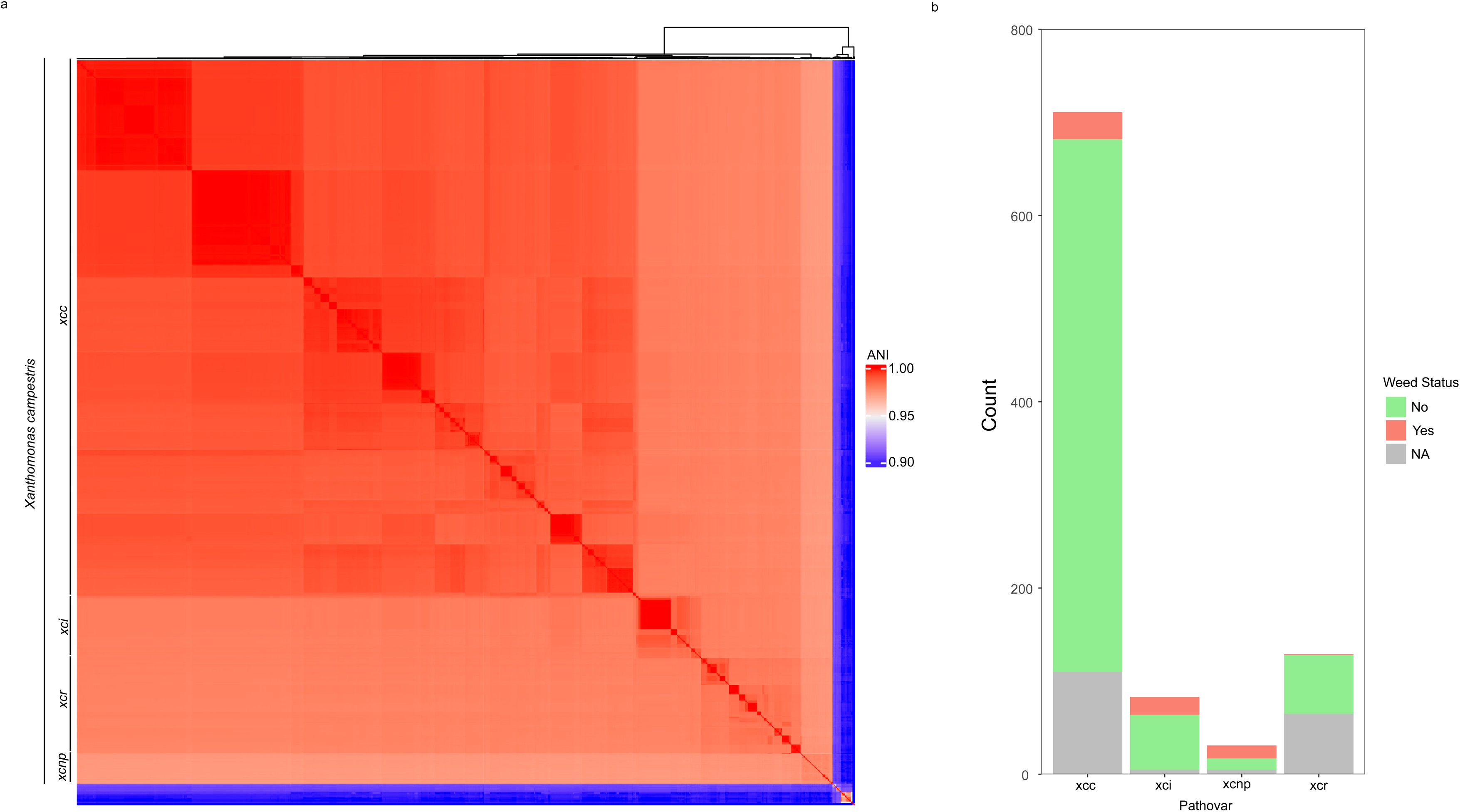
Initial genome validation and host isolation source mining. A) Heatmap representing all isolates of *Xanthomonas campestris* (N = 1003) on 4/18/2026 using average nucleotide identity (ANI) after mapping to type strain ATCC-33913 of *X. campestris*. Pathovars of *X. campestris* are labeled on the left and strains below the ani cut-off of 0.95 were removed from the study. B) Counts of isolation sources, determining the number of weed-isolated strains. All major pathovars of *X. campestris*, with the exception of *xcr* were found to have sizable numbers of strains isolated from weedy species.

**Supplementary Figure 2.**
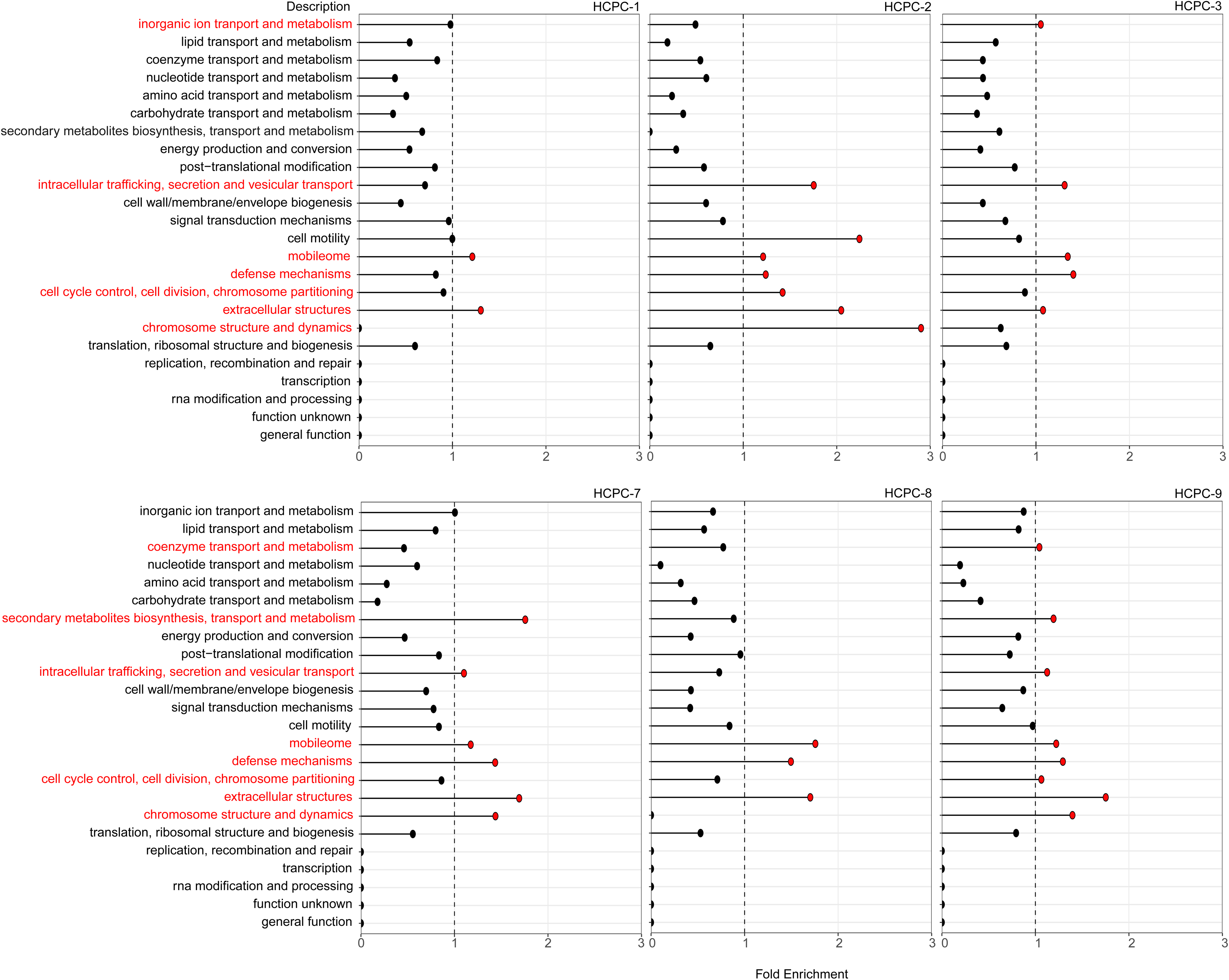
Location and density analysis analysis of correlated genes. Correlated genes (vtest > 2) were mapped onto the representative genomes of each HCPC, with the exception of HCPC-4 which had a highly fragmented genome. Strains include A) HCPC-1 (B100), B) HCPC-2 (22-193), C) HCPC-3 (ATCC-33913), D) HCPC-5 (XUMN), E) HCPC-6 (11-18), F) HCPC-7 (25-4), G) HCPC-8 (21-089), H) HCPC-9 (CFBP-1606). Clusters HCPC-2 and HCPC-9 both have associated plasmids that are visualized along with the main chromosome. The vtest score was used to color each gene, with brighter colors representing strong vtest values. A gene count area plot, where the y-axis represents raw count of genes per 1 kb, is included to better visualize density of genes in the crowded genome maps.

**Supplementary Figure 3.**
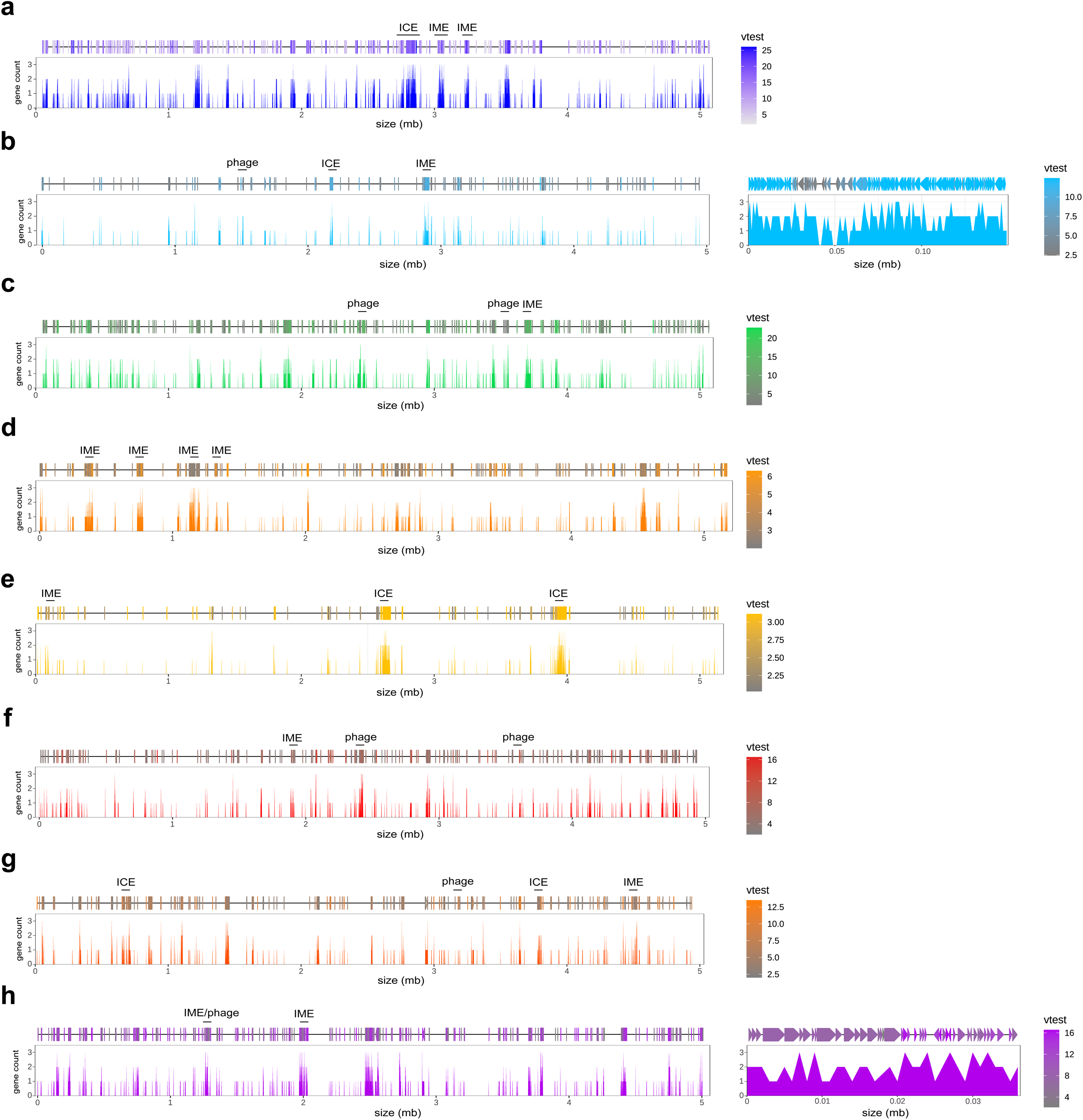
Clustering of plasmid sequences from high-quality genomes. Plasmids are grouped using a Jaccard distance cut-off of 0.8 to cluster similar plasmids. Two plasmids failed to meet this cut off and were labeled as pRare. The BAPS cluster of the corresponding strain is displayed in the heatmap.

**Supplementary Figure 4.**
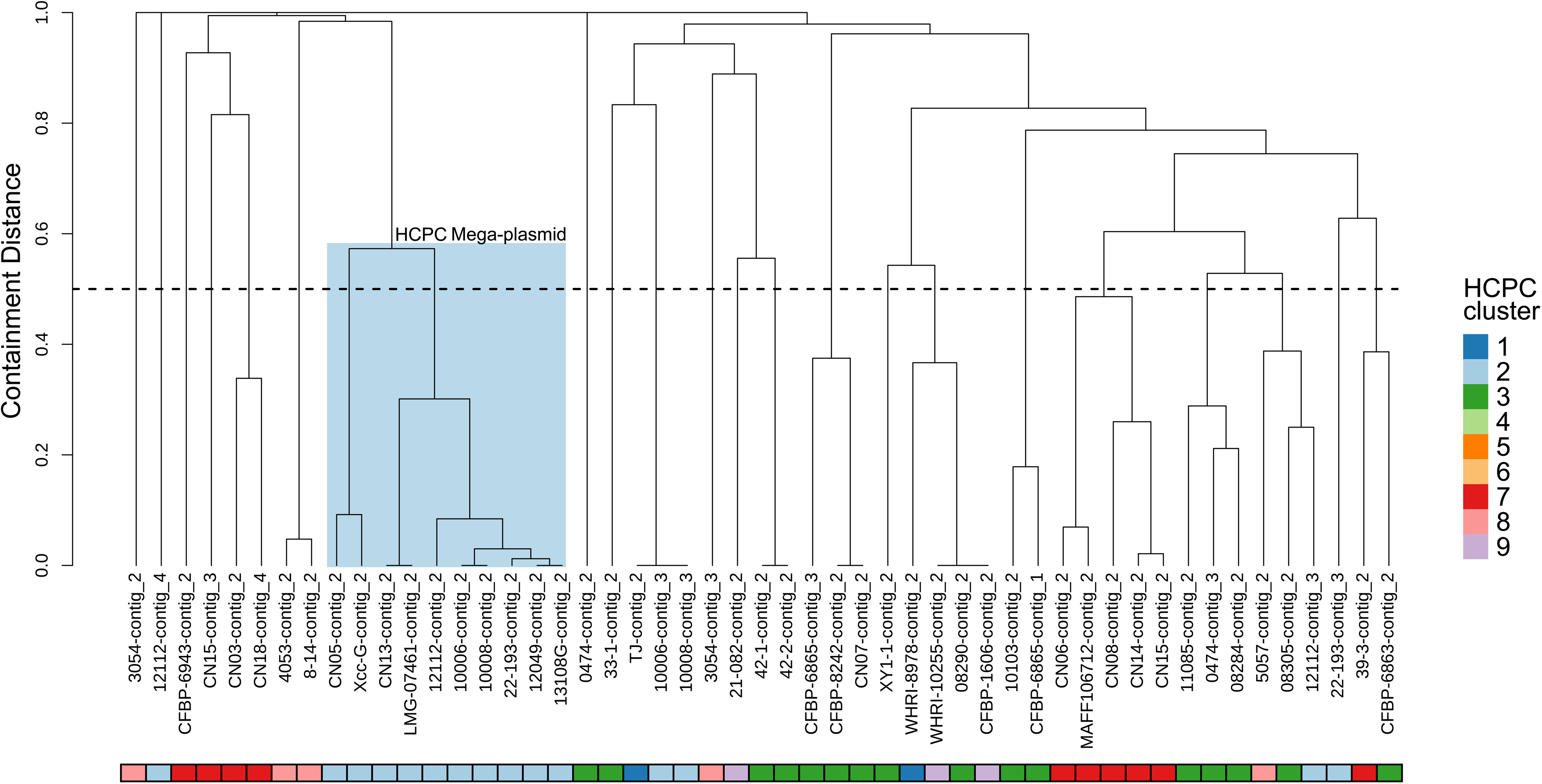
Clustering of predicted ICE elements locates a conserved element across *Xcc*. A) Global analysis of all predicted ICE and IME elements within the pangenome. The type of element (ICE or IME) and the BAPS cluster it was isolated from are denoted in heatmaps 1 and 2. B) Extraction of ICE1 (ICE-avr) in panel a.

**Supplementary Figure 5.**
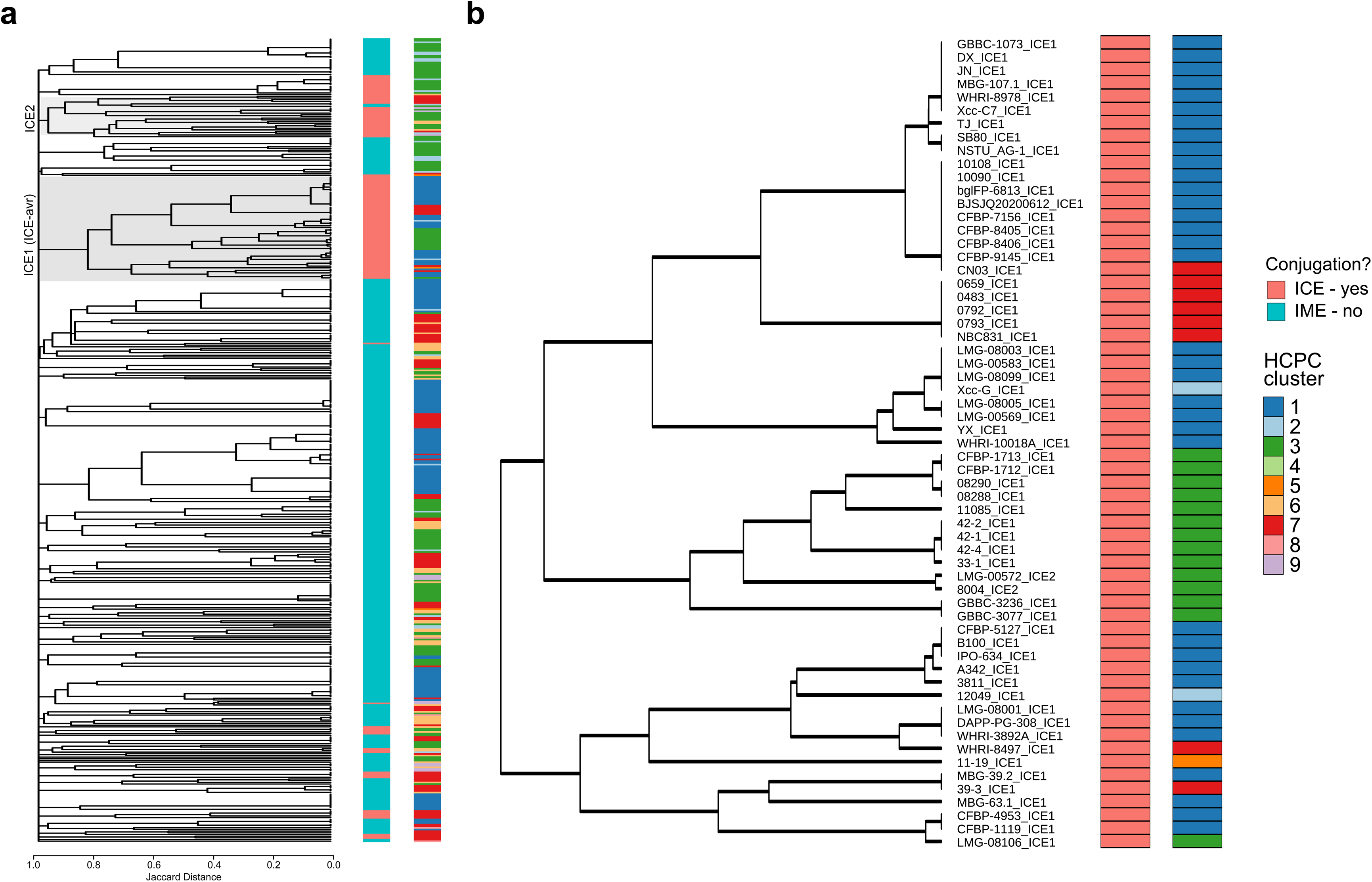
Multiple sequence analysis of unique *avrB* sequences reveals multiple truncation events. Unique protein sequences were collected from all proteins in the avrB gene family. The count for each unique isoform is provided next to the name. Reading frames which are predicted to be truncated and likely non-functional are italicized.

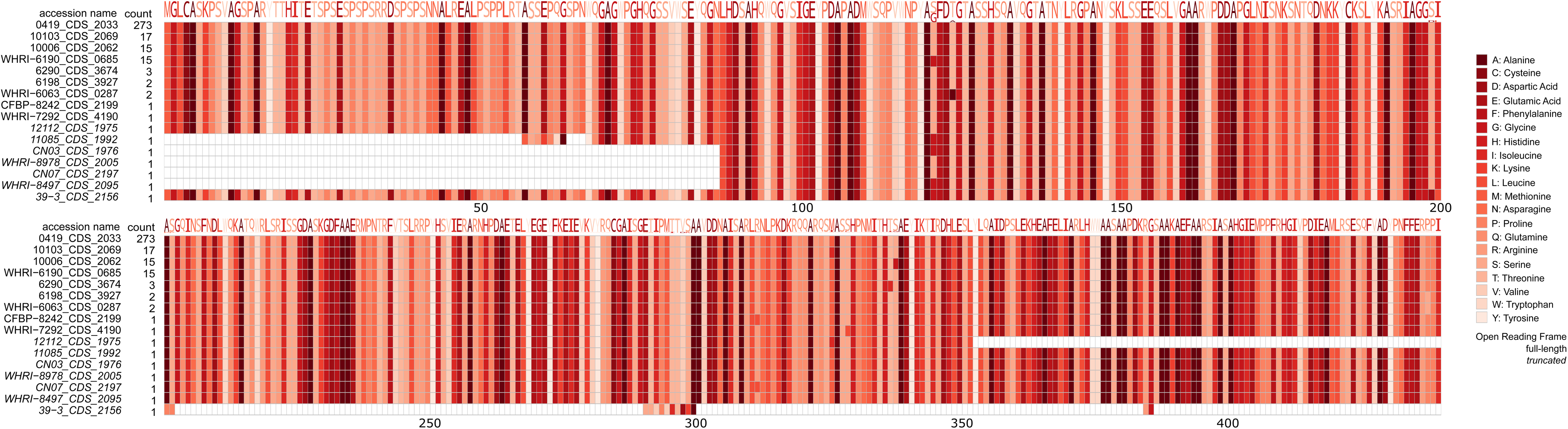

## Notes

### Competing Interest Statement

The authors have declared no competing interest.

